# Enumeration of Citrus endophytic bacterial communities based on illumine metagenomics technique

**DOI:** 10.1101/2022.01.13.476241

**Authors:** Sehrish Mushtaq, Muhammad Shafiq, Tehseen Ashraf, Muhammad Saleem Haider, Sagheer Atta

**Author notes:** These authors contributed equally to this work. These authors also contributed equally to this work.

## Abstract

Citrus is a valuable crop in Pakistan because it is rich in vitamin C and antioxidants. Huanglongbing (HLB) has an influence on citrus production around the world caused by a bacterium *“Candidatus liberibacter asiaticus”* (CLas), africanus and americanus. The structure and diversity of bacterial species in various ecosystems can be quickly examined using NGS. This approach is considerably quicker and more precise than outdated methods. Healthy or citrus greening infected leaf samples of Grapefruit, *Citrus aurantifolia*, and *Citrus reticulata* Blanco was used for diversity analysis. In this study high throughput, NGS technique was used to access the population of both cultivable and non-cultivable bacterial endophytes from citrus leaves, by using PCR amplicons of 16S rDNA sequences (V5–V7 regions) with Illumina Hi seq. As a result, a total number of 68,722 sequences were produced from the test samples. According to the NGS-based diversity classification, the most common genera of exploited bacterial endophytes were Proteobacteria, Firmicutes, Bacteroides, Cyanobacteria, and Actinobacteria. *Citrus aurantifolia* and *Citrus paradisi* showed almost equal diversity, whereas *Citrus reticulata* Blanco had a higher proportion of Proteobacteria and Cyanobacteria in their leaves. To determine alpha diversity (AD), additional data was analyzed using statistical indices such as Shannon, Chao1, and Simpson. According to the inverse Simpson diversity index, the abundance of the microbial population in six different citrus samples was 0.48, 0.567, and 0.163, respectively. The metagenomics of microbiota in plant tissues was successfully recorded by NGS technology, which can help us learn more about the interactions between plants and microbes. This research is the first step toward a better understanding of 16SrRNA-based metagenomics from citrus in Pakistan using Illumina (Hi seq) Technology.

## Introduction

Pakistan is one of the world’s largest citrus producers, ranking 13^th^ in total citrus production. Citrus is highly important due to its economic and nutritional benefits. Kinnow is a useful fruit that occupies the first place among all fruits in terms of both area and production [1]. The total area under citrus cultivation during 2014-15 was 192832 hectares with a production of 2395550 (tons) [2]. Punjab is home to nearly all of the world’s citrus groves. With more than 75% production of total citrus fruits, 29.55% of the total area is planted in citrus and 60% in kinnow. About 90% of all citrus exports are kinnow. Major Citrus species cultivated in Pakistan are as follows; Grapefruit, Sweet orange, Mandarin, Lemon Lime, Bitter orange [3].

Citrus diseases have emerged as a possible threat to global citrus productivity. HLB, a disease caused by three gram-negative, phloem-limited alphaproteobacteria: *“Candidatus liberibacter asiaticus”* (CLas), africanus, and americanus have a major effect on citrus production worldwide [4]. However, different CLas strains have been recorded from the United States, specifically from Florida [5–8], Iran [9], Mexico [10], Australia [11], and Pakistan [12]. HLB is distinguished by less nutrient transfer, resulting in a variety of distinct effects, including yellow shoots, branch dieback, green fruit remaining, lopsided fruit, reduced size and eventually tree death [13]. The plant microbiome plays a part in different aspects of plant health and disease, including growth rate, vigor, and tolerance, inflammation, and disease resistance [14, 15]. Understanding how the microbiome affects and communicates with the plant would entail the application of several experimental methods, including a meta-analysis of broad Meta datasets with critical variables relevant to plant health, protection, and disease [16].

NGS is a culture-independent method that is useful for the study of the entire microbial population within a sample. High-throughput sequencing technologies [17] refer to a group of tools that can be used to sequence DNA of various base pairs faster and cheaper than previous methods. NGS sequence of DNA fragment (16S rRNA) in the form of reading (short DNA fragment) as compared to reference sequences from databases in lesser time to identify the related bacterium with this fragment [18, 19]. There are various studies of 16SrRNA gene base sequencing for targeted amplification of bacterial communities [20]. Although, in this era of science researchers are using the most effective variable (V) region of the 16SrRNA gene for sequencing, with many studies selecting to examine more than one region as no single region has been shown to optimally differentiate among bacteria [21, 22]. All nine Variable regions of 16S rRNA displayed bacterial diversity and the most important step is determining which variable region to sequence, since classification bias variable region has been found previously [23]. The use of PCR-based molecular techniques (polymerase chain reaction) has made it possible to research the total diversity of microbes in the natural environment without the cultivation of microbes [24]. These new advanced techniques are valuable in increasing our understanding of the microbial communities regardless of some amplification biases demonstrated due to the selection of suitable primers, the concentration of template, and the number of amplification cycles [25, 26].

NGS-based microbial community research has paved the way for the development of novel culture-independent bacterial strains capable of identifying biological control agents against the HLB pathogen (*Candidatus liberibacter asiaticus*). The study of biological control organisms’ natural microbial niches, which are close to those of pathogens, could lead to more successful disease control. Microbial diversity associated with citrus leaf (phloem) can be identified by either cultivation-dependent or cultivation-independent methods. On the other hand, the fraction of bacterial diversity measured using previous culture techniques accounts for just 0.1 to 10% of the overall estimated diversity [27, 28], suggesting that laboratory culture techniques are substantially biased. However, it is a fact that the majority of dominant bacteria present in environmental samples are uncultivable [29–32]. 16S rDNA-based phylogenetic analysis has been widely used to classify microbial diversity in different environmental niches, such as soil [30], plants [33, 34], subsurface sediments, and rocks [35]. The primary aim of this research was to determine whether bacteria other than *Ca. Liberibacter spp*. is associated with the citrus greening disease.

Microbial diversity research is important for recognizing the microbial flora that exists on plants in their natural environment. The diversity of bacterial endophytes from citrus in Pakistan is the focus of this report, which is based on preliminary research. The uncultivable and cultivable fraction of bacteria is first time exploited from citrus leaves through the Illumina metagenomics technique (Hi seq) in Pakistan. There has been an increased recognition that it is necessary to pay more attention to this area. NGS (next-generation sequencing) is an incredibly valuable technique to access the uncultivable fraction of bacterial endophytes in plant tissues. This could help us better understand the microbes that live on plant surfaces in natural conditions and how they interact. To the best of our understanding, this is Pakistan’s initial 16SrRNA-based metagenomics study from citrus leaves using Illumina (Hi seq). The main objectives of this research were to investigate the microbial species associated with the leaf midribs of HLB symptomatic and asymptomatic citrus (*Citrus aurantifolia*, *Citrus paradisi*, *Citrus reticulata* Blanco) trees and also to know their relative abundance, and phylogenetic diversity by using high-throughput 16S rDNA (V5-V7) next-generation sequencing through Illumina (Hi-seq).

## Materials and Methods

### Samples collection and DNA isolation

Leaf samples (healthy/infected) of grapefruit, *Citrus aurantifolia*, and *Citrus reticulata* Blanco were obtained from IAGS, Pu, Lahore backfields and preserved at −80°C. Citrus plants that were six years old were used for this experiment and five leaves per plant were taken as a sample and stored at −80°C. To extract soil particles, every plant leaf was washed and cleaned under running tap water. The leaves were washed in autoclaved water with a few drops of Tween-20 and set aside to drain for 10– 15 minutes. Then they were cut into 4–5 bits, each measuring 2–3cm in length. Surface sterilization was carried out using the methods defined by [36], with some variations in the Ethanol conc. and sterilization time. Soft tissue were submerged in ninety percent ethanol soln. for 5 minutes, then in a 3 percent sodium hypochlorite solution for 2 minutes, and finally in 75% ethanol (3 min). The disinfected leaves were drained in a laminar flow hood after being rinsed three times with autoclaved distilled water. The surface-sterilized tissues (control) and the last rinsing water were inoculated onto nutrient agar plates to confirm the efficacy of the surface sterilization procedure. Any bacteria growth in the control agar plates within 24 hours of incubation (30°C±2°C) indicates ineffective surface sterilization. The complete genome of DNA was extracted using the CTAB method (cetyl trimethyl ammonium bromide), as defined by [37]. At A260/280 nm (1.9–2.0), the isolated DNA was quantified and tested for purity (Nanodrop at School of Biological Sciences Pu, Lahore) and stored at −20°C before being processed. For NGS (Illumina Hi seq), these quantified DNA samples were sent to the Novo gene (leading-edge genomics services and solutions).

### Generation of Amplicon

The bacterial genomic DNA concentration in leaf tissue samples was normalized to 10 ng/L. The conserved regions of 16S rRNA were amplified using PCR (V5-V7-WBI-NV2018010942). Phusion^®^ High-Fidelity PCR Master Mix was used to prepare the PCR library (New England Biolabs). Briefly, 25 μL PCR reaction comprises DNA (6 μL), 12.5 μL of (2x) Master KAPA Hi-Fidelity DNA polymerase (1 U), primer (10 μM) 1.5 μL (each), and distilled autoclaved water. PCR reactions were initiated with 95°C for 3 min (denaturation cycle) followed by 24 cycles at 98°C for 20 sec, 55°C for 15 sec, and 72°C for 10 sec, and ended at 72°C for 1 min (Extension step). Mix the same amount of 1X loading buffer (with SYB green) with PCR products and run electrophoresis on 2% agarose gel for detection. Samples with a bright main strip between 400-450bp were selected for further studies. PCR products were Gel purified by using Qiagen Kit Manufactured by (Qiagen, Germany).

### Library preparation and sequencing

Sequence libraries were generated using the TruSeq^®^ DNA PCR-Free Sample Preparation Kit (Illumina, USA) following the instructions given. The quality of the library was analyzed by the Qubit@ 2.0 Fluorometer (Thermo Scientific) and the Agilent Bioanalyzer 2100 system. To conclude, the library was sequenced on the Illumina HiSeq 2500 platform and 250 bp paired-end reads were produced. A preliminary study of the illustration and base call was performed on the HiSeq instrument. Hi Seq (ultra-high-throughput) was used to de-multiplex data and exclude reads in FASTQ format that failed the Illumina purity filter (PF = 0). The forward and reverse reads of raw data were combined using the mother pipeline alignment method. Following that, they were trimmed and filtered by deleting the bases with rating scores less than or equal to 2, the maximum number of N accepted = 4, the maximum number of homopolymers accepted = 8, and the contaminant removed. All tests were performed using the Mothur pipeline program software (http://www.mothur.org/wiki/).

### Classification of bacteria

The SILVA rRNA database and the Silva database were used to assign operational taxonomic units (OTUs) were assigned to the retrieved read sequences produced from the leaf samples. We used the mother pipeline’s “splitting by classification “process to assign OTU.

### Statistical analysis

All of the data was processed using one-way ANOVA. The Statistical Package for Social Science (SPSS) was used to conduct the analysis, Tukey’s Studentized Range Test HSD (0.05) has been used to compare the means, and p values less than 0.05 which were considered statistically significant.

### Diversity Analysis

Alpha and beta are two methods of diversity analysis that are commonly used to find diversity using NGS. Alpha diversity (AD) is used to analyze the complexity of species diversity in the experiment by diversity indices, including Observed-Species, Chao1, Shannon, Simpson, ACE, and Good-coverage. All of these indices were measured with QIIME and viewed with the R program. Beta Diversity (BD) Analysis was used to assess differences in sample species complexity. Beta Diversity was measured using QIIME software Unit fraction metrics (unifrac), as weighted and unweighted. Unifrac is a method of calculating the phylogenetic distance between taxonomic groups in a tree as a percentage of the length of the branch that contributes to ancestors from either one or both origins. Arithmetic Means in an Unweighted Pair-group Method (UPGMA) QIIME was used to perform clustering, a hierarchical clustering technique that uses average linkage to interpret the distance matrix.

## RESULTS

### DNA extraction for Next-Generation Sequencing (Hi seq)

Leaf samples of both healthy and symptomatic *Citrus paradisi*, *C. reticulata* Blanco, and *C. aurantifolia* were obtained from the backyard of the Institute of Agricultural Sciences and preserved at-80°C. The CTAB method was used to separate DNA from the leaf samples. To access the diversity of cultivable vs. non-cultivable bacteria, isolated DNA was electrophoresed in a 1% agarose gel (to verify DNA) and, quantified through nanodrop before further processing. sent for Illumina Hi seq NGS technique.

### Sequencing and data processing

The Illumina paired-end network was used to sequence the PCR amplicon yielding raw reads (Raw PE) with paired ends of 250 bp that were then extracted and Clean Tags were obtained after being pretreated. To obtain Effective Tags, Clean Tags that included chimeric sequences were identified and excluded. The data output indicates data interpretation and QC status (Table 1).

**Table 1:** Data processing and QC (quality control) stats of citrus (*Citrus aurantifolia*, *Citrus paradisi*, *Citrus reticulata* Blanco) samples.

| Sample abbreviations used in this study | Raw PE (#) | Raw Tags (#) | Clean Tags (#) | Effective Tags (#) | Base (nt) | AvgLen (nt) | Q20 | Q30 | GC % | Effective % |
| --- | --- | --- | --- | --- | --- | --- | --- | --- | --- | --- |
| *C.aurantifolia* Healthy (SM1) | 42,914 | 41,860 | 39,641 | 26,813 | 9,982,254 | 372 | 98.60 | 97.08 | 53.71 | 62.48 |
| *C. aurantifolia* Infected (SM2) | 35,655 | 34,844 | 33,190 | 22,880 | 8,21,549 | 372 | 98.66 | 97.22 | 53.77 | 64.17 |
| *C. paradisi* Healthy (MK1) | 68,722 | 67,168 | 63,631 | 55,999 | 20,820,815 | 372 | 98.58 | 97.05 | 53.36 | 81.49 |
| *C. paradisi* Infected (MK2) | 57,447 | 56,216 | 53,447 | 47,782 | 17,789,958 | 372 | 98.58 | 97.07 | 53.57 | 83.18 |
| *C. reticulata* Blanco Healthy (MA1) | 33,837 | 33,037 | 31,226 | 24,496 | 9,149,614 | 374 | 98.53 | 96.98 | 55.43 | 72.39 |
| *C.reticulata* Blanco Infected (MA2) | 30,903 | 30,223 | 28,359 | 25,189 | 9,440,842 | 375 | 98.45 | 96.82 | 55.99 | 81.51 |

### OTU Clustering and species annotation

All Successful Tags were grouped into OTUs based on 97% DNA sequence similarity to evaluate the species diversity in each sample. Detailed information gathered from a variety of samples, such as Tag annotation data, effective Tags data, and low-frequency Tags data was collected during the construction of OTUs. The statistical data set is organized as follows (Fig. 1).

**Figure 1:**
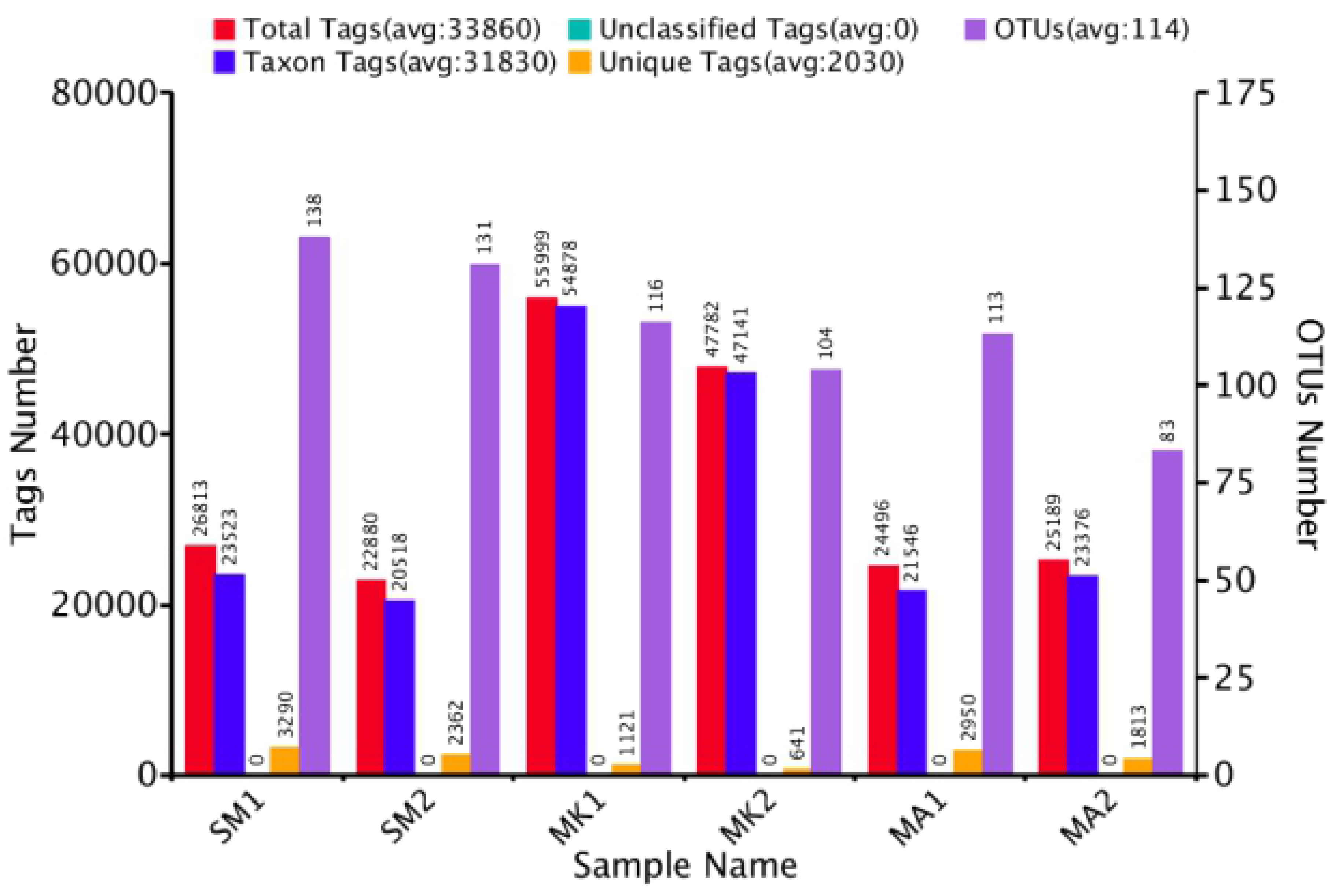
Statistical analysis of the tags and operational taxonomic units of each tested citrus leaf sample

### Phylogenetic Tree

R and D software was used to select independently the most common top ten genera of specific species with high relative abundance by default) for the construction of a phylogenetic tree [51]. Actinobacteria, Cyanobacteria, and Firmicutes, as well as Proteobacteria, were identified that belong to the phylum (Alpha, beta, and gamma). The research samples were found to be infected with eight orders and nine groups of bacteria (figure tree of particular species in samples SM-1/SM-2 (Asymptomatic/Symptomatic *Citrus aurantifolia*), Mk-1/MK-2 (Asymptomatic/ Symptomatic *Citrus paradisi*); MA1/MA2 (Asymptomatic/Symptomatic *Citrus reticulata* Blanco). In this diagram, the four major phyla are represented (Fig. 2).

**Figure 2:**
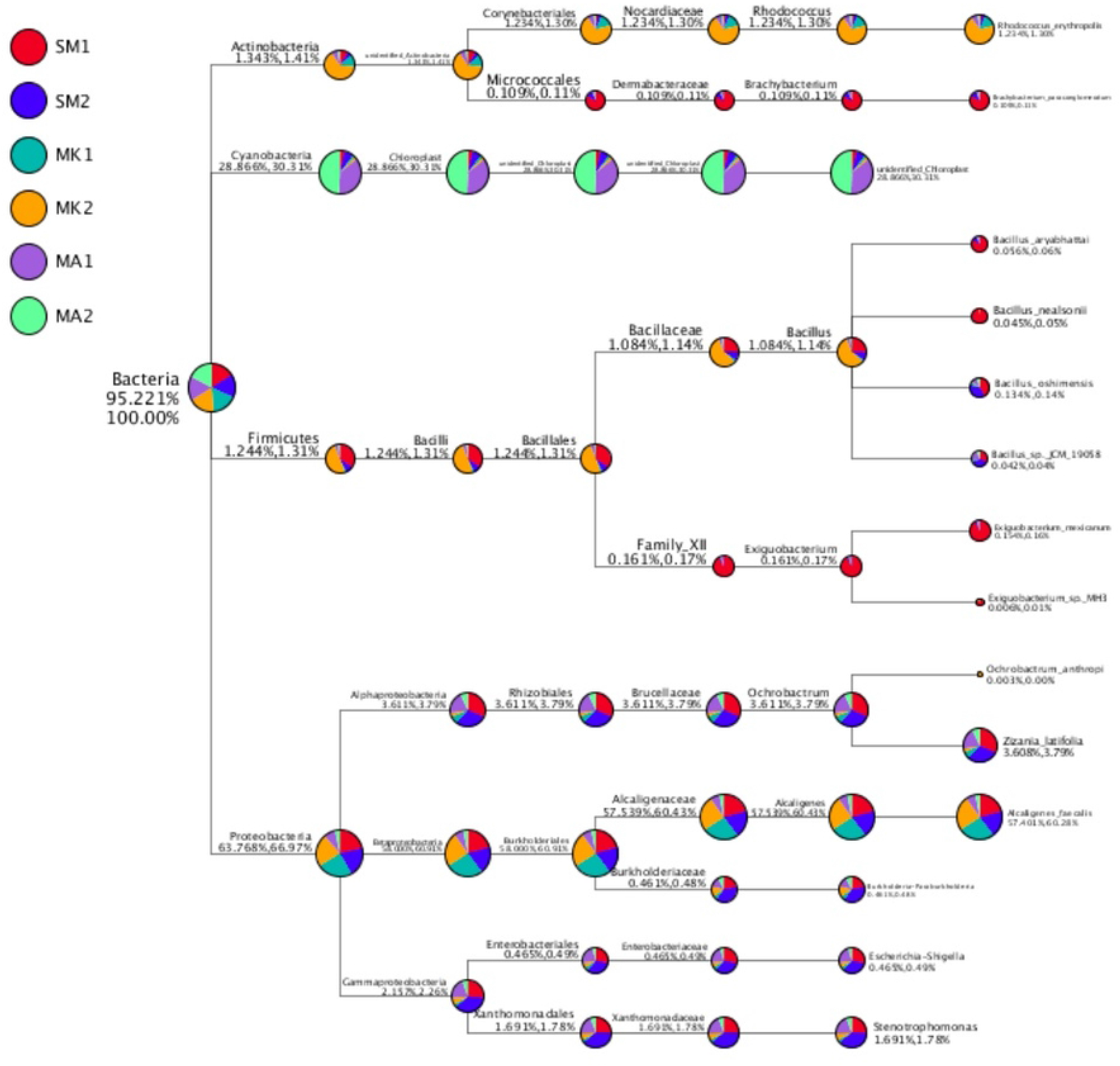
Taxonomy tree of specific species in citrus leaf samples.

### Relative Abundance of Species

To structure the scattering of relative abundance of species in histograms, the top 10 species in each taxonomic rank were chosen. The distribution of the phyla can be seen in (Fig. 3) and the relative abundance of bacterial species in normal vs. infected leaves revealed that SM1/SM2 (*Citrus aurantifolia* asymptomatic/symptomatic) has a higher proportion of Proteobacteria, whereas the infected one has a smaller proportion of other phyla, with Cyanobacteria dominating among them. MKI/MK2 (Citrus paradisi asymptomatic /symptomatic) showed a similar pattern. MA1/MA2 (*Citrus reticulata* Blanco asymptomatic/ symptomatic) had 40% Proteobacteria and 60% Cyanobacteria, while MA2 had 20% Proteobacteria and the remaining 80% Cyanobacteria and another phylum. The relative abundance of bacteria is calculated by integrating both symptomatic and asymptomatic bacteria into one group were represented through; Bac-1 (Citrus aurantifolia) community revealed 90% Proteobacteria and just around 10% cyanobacteria. While in Bac-2 (*Citrus paradisi*) group only Proteobacteria was found in abundance. Bac-3 (Citrus reticulata Blanco) community, on the other hand, had a 25% proportion of Proteobacteria and a 75% proportion of Cyanobacteria and others.

**Figure 3:**
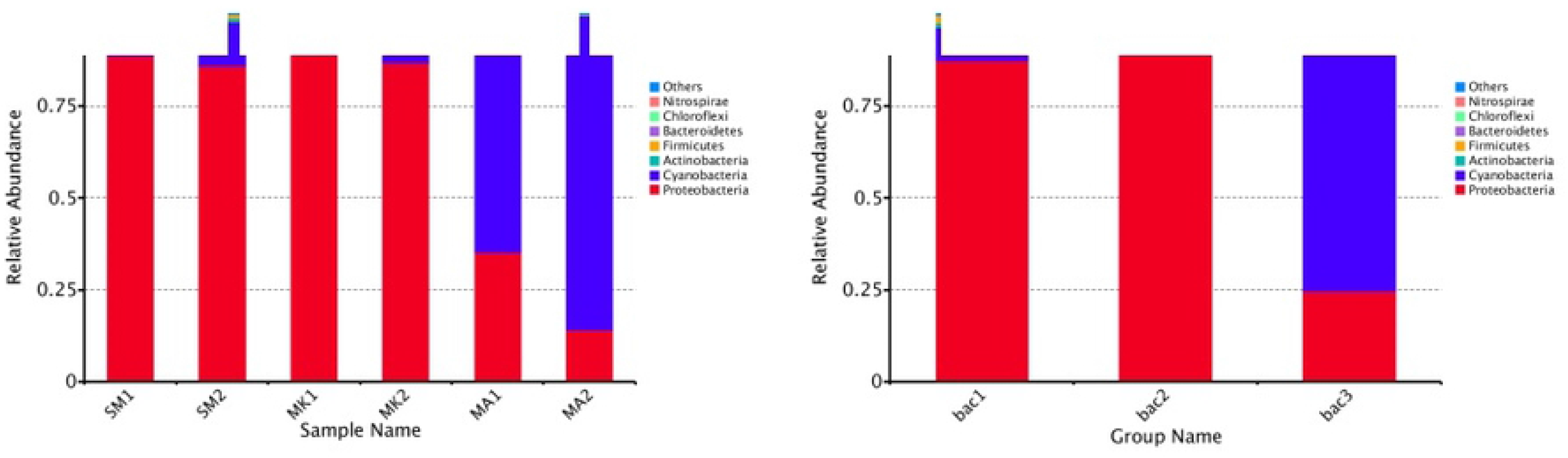
Relative abundance of bacterial Species at phylum level from citrus leaves

### The Phylogenetic tree

The top hundred taxa have been selected, and the evolutionary tree was built by aligning the sequences. Each genus’ relative abundance was measured as shown in (Fig. 4).

**Figure 4:**
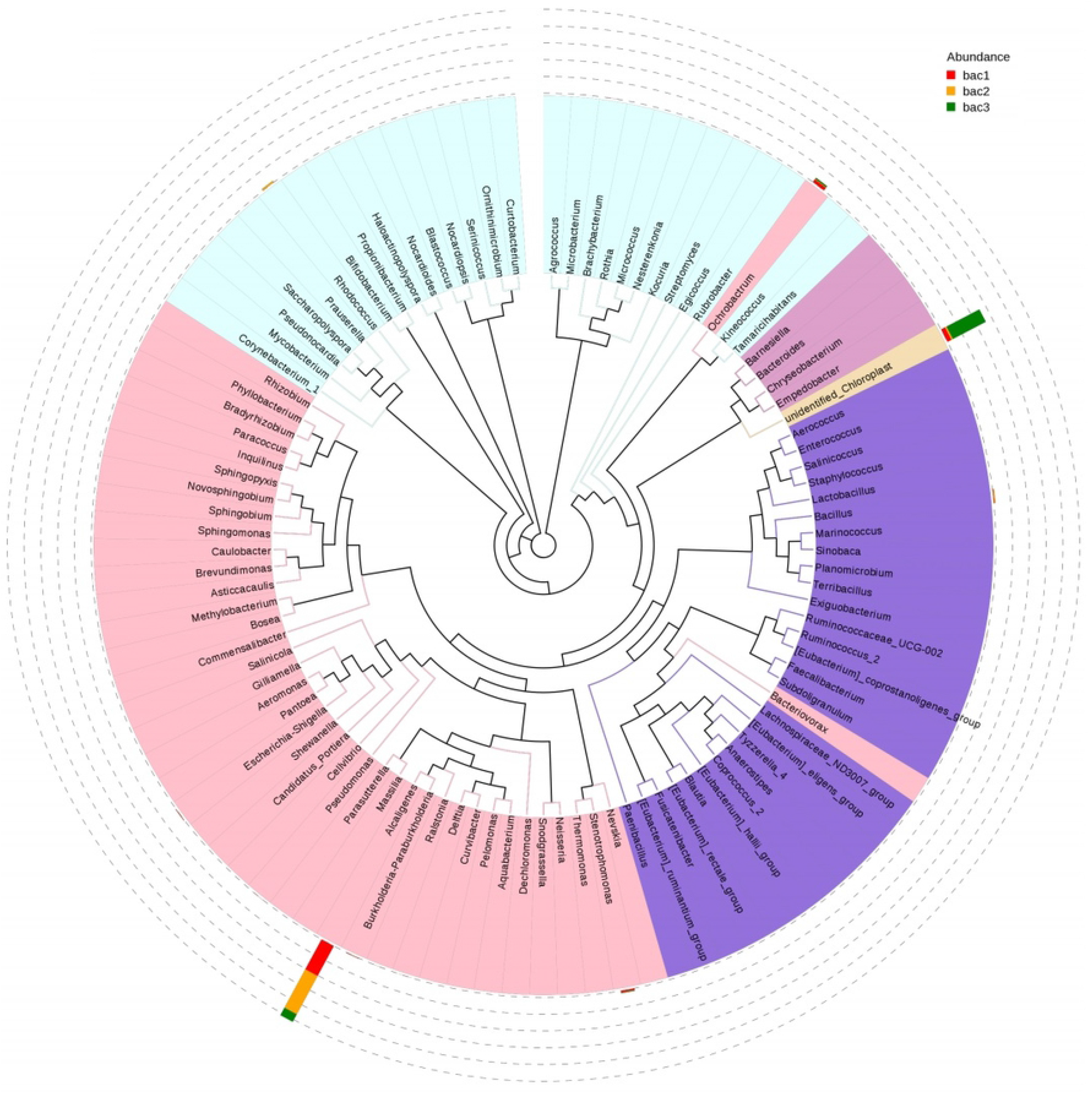
The evolutionary tree based on the genus of Bacterial endophytes from citrus leave

Venn diagrams were also constructed based on operational taxonomic units of the identified bacteria from citrus leaf samples as shown in (Fig.5).

**Figure 5:**
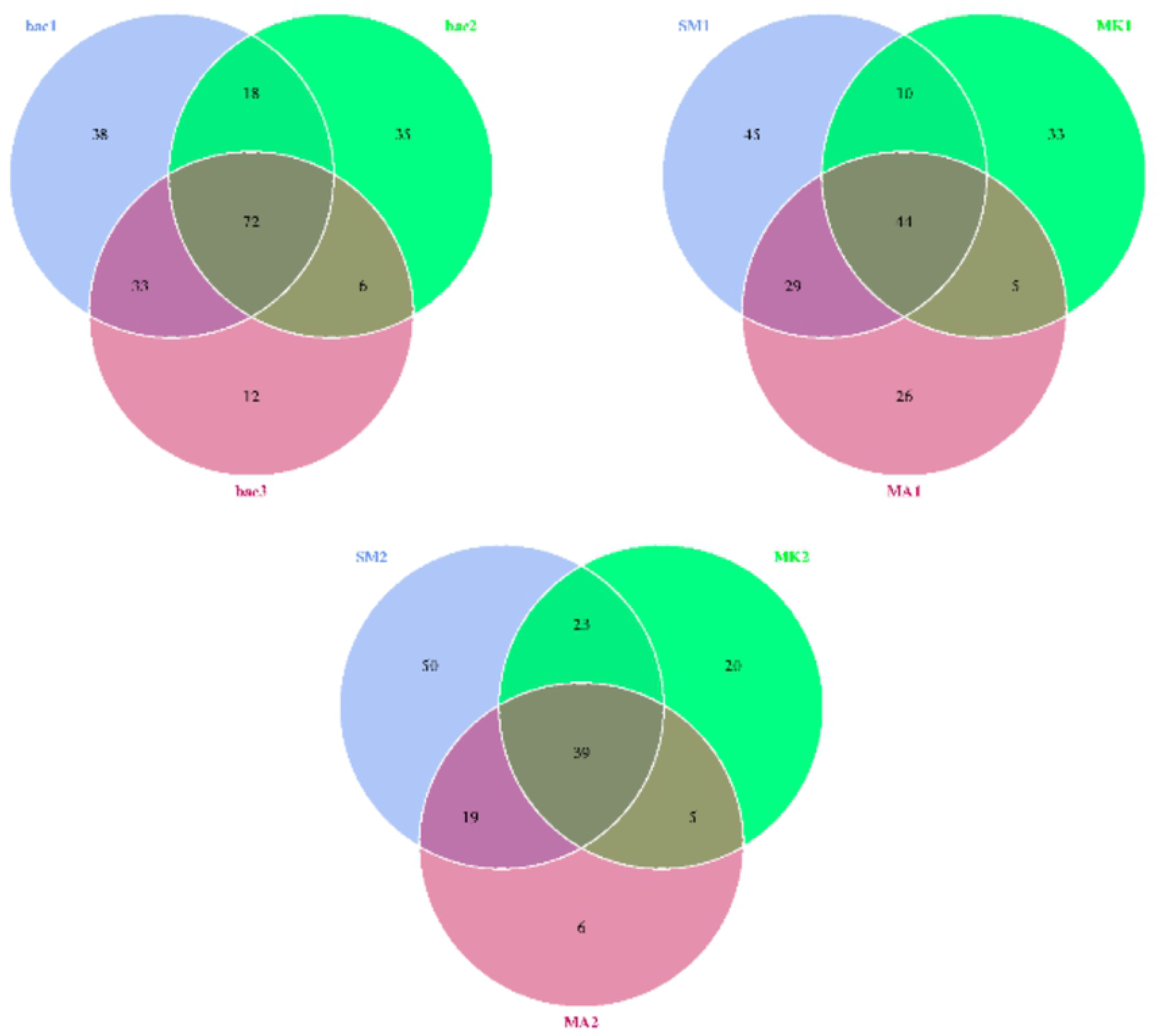
Venn diagram constructed based on operational taxonomic units of the bacterial diversity from citrus leaf samples

### Alpha and Beta diversity Analysis

OTUs with 97% sequence identity are assumed to be homologous among species and statistical indices of AD are listed in (Table 2).

**Table 2:** Statistical analysis of alpha diversity (AD) indices from NGS data of citrus leaves.

| *Sample Abbreviations used in this study* | **No of species observed** | **Simpson** | **Shannon** | **Chao1** | **ACE** | **Good coverage** | **PD whole tree** |
| --- | --- | --- | --- | --- | --- | --- | --- |
| *C.aurantifolia* Healthy (SM1) | 128 | 0.484 | 2.120 | 141.571 | 138.761 | 0.999 | 9.691 |
| *C.aurantifolia* Infected (SM2) | 131 | 0.567 | 2.290 | 138.241 | 144.296 | 0.999 | 9.428 |
| *C.paradisi* Healthy (MK1) | 92 | 0.163 | 0.751 | 113.136 | 130.639 | 0.998 | 8.407 |
| *C.paradisi* Infected (MK2) | 87 | 0.307 | 1.245 | 102.812 | 105.703 | 0.999 | 8.125 |
| *C.reticulata* Blanco Healthy (MA1) | 104 | 0.741 | 2.419 | 120.714 | 130.361 | 0.999 | 8.699 |
| *C.reticulata* Blanco Infected (MA2) | 69 | 0.539 | 1.602 | 84.833 | 88.361 | 0.999 | 6.379 |

### Beta Diversity Indices and heat map

Unweighted vs. Weighted Unifrac distances, which are phylogenetic indicators that are commonly used in current bacterial community sequencing projects, were chosen to quantify the dissimilarity coefficient between pairwise samples. In this graph, a heat map centered on the weighted vs. unweighted Unifrac distances is plotted (Fig.6). The red section of the triangle suggests that there is less beta variety among samples, whereas the yellow portion indicates that there is more beta diversity among samples (SM2, MK1, MK2, and MA1).

**Figure 6:**
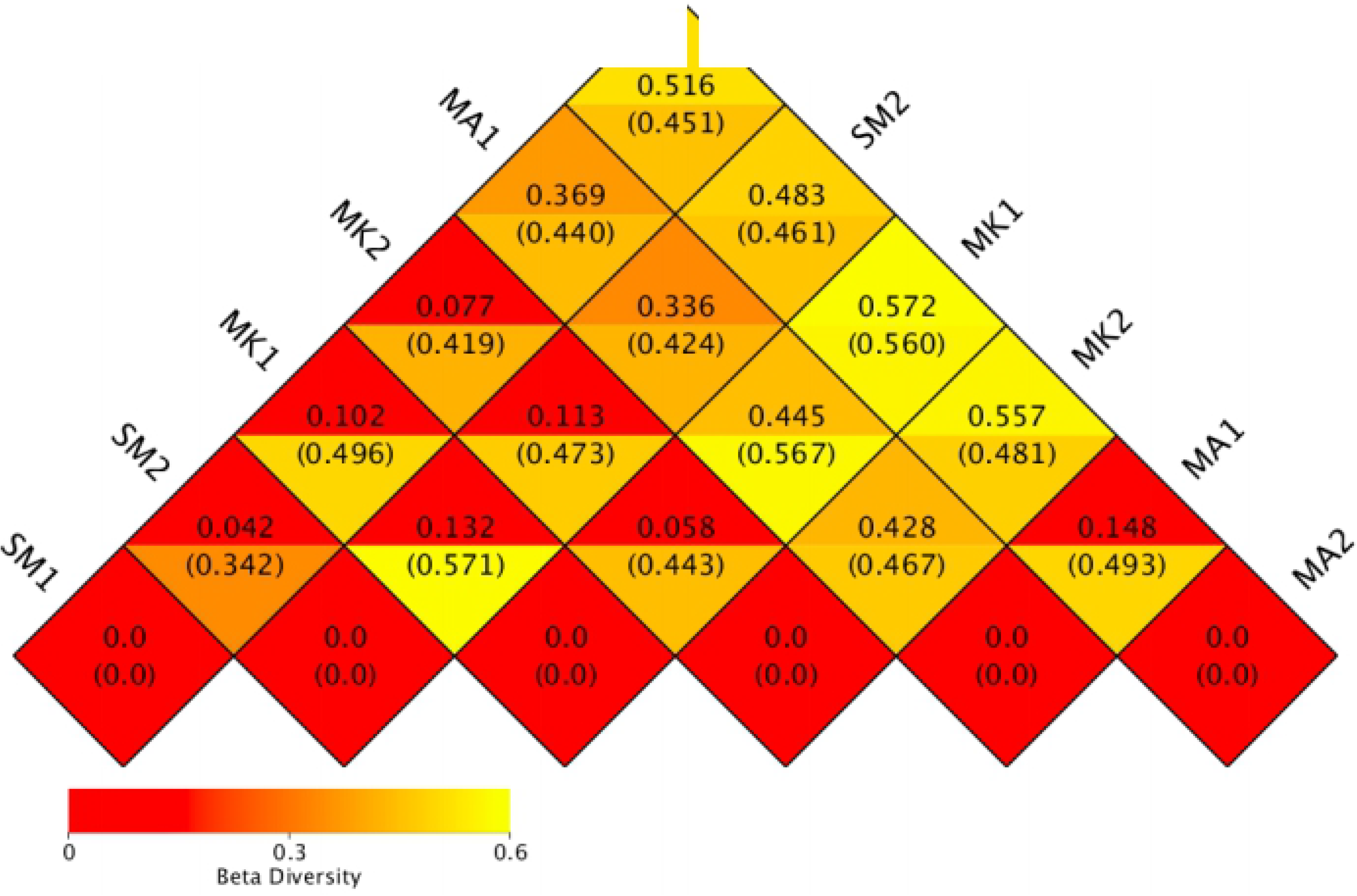
Illustrates beta diversity analysis (Heat map) based on Weighted/Unweighted Unifrac distances.

### Unweighted Pair-group Method with Arithmetic Mean (UPGMA)

Clustering analysis and the construction of a clustering tree were used to investigate the similarities between different samples. The (UPGMA) procedure with arithmetic mean is a type of hierarchical clustering method used for classifying ecosystem samples. The following are fundamental concepts of UPGMA methods. The samples with the shortest distance were being grouped, and then a new sample is generated. It has a branching point in the middle of the two initial samples. After computing the average distance between the newly created “sample” and other samples, the closest two samples can be used to repeat the procedures adopted earlier in this section. Until all of the samples are clustered together, a complete clustering tree can be obtained. Before conducting UPGMA cluster analysis, the Weighted Unifrac distance matrix and the Unweighted Unifrac distance matrix were calculated. They could be seen in a graph that included the clustering results as well as every sample’s phylum-specific relative abundance (Fig. 7a and 7b).

**Figure 7:**
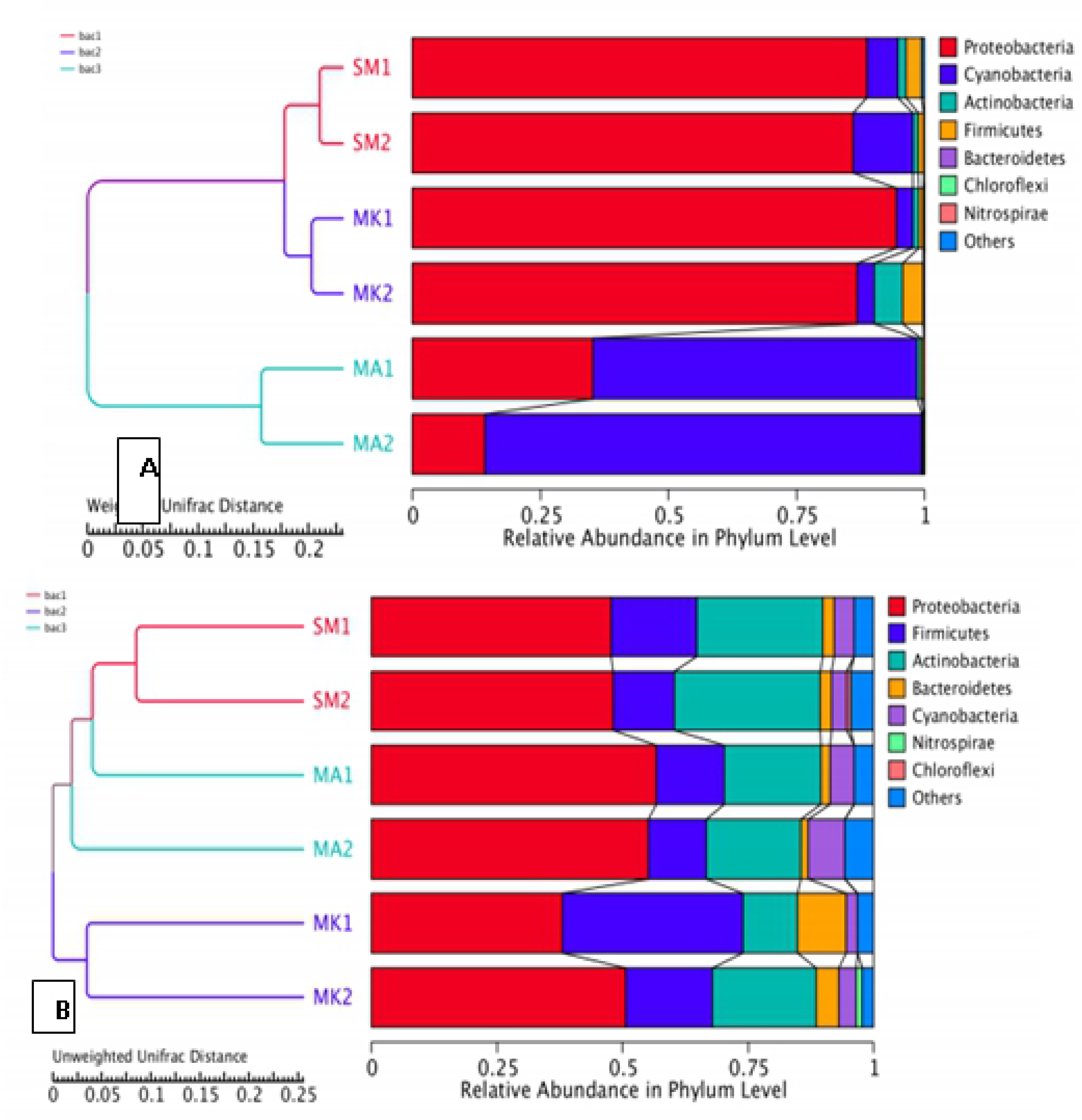
UPGMA cluster tree based on a) Weighted Unifrac distance b) Un Weighted Unifrac distance showing the relative abundance of bacterial species at phyla level.

The SM1/SM2 and MK1/MK2 clusters in the same clade had more or less similar bacterial diversity, according to the UPGMA cluster tree based on the Weighted Unifrac distance tree, however, MA1/MA2 displayed a distinct configuration and is in a different clade, indicating that *Citrus reticulata* Blanco has a different bacterial diversity than *Citrus aurantifolia* and *Citrus paradisi*. The UPGMA cluster tree based on unweighted unifrac distance displays a variable pattern if compared to the weighed unifrac distance tree. MK1/MK2 is in a distinct clade in unweighted unifrac distance trees, whereas the other two groups are all in the same clade, as seen in (Fig.7b). MK1/MK2 had a higher proportion of Bacteroides than the others. As a whole, the most common genera found in three samples were Proteobacteria, Cyanobacteria, and Actinobacteria.

## DISCUSSION

A microbial community study is a fast way to learn regarding the structure and functioning of bacterial communities, and it could contribute to the isolation and detection of new bacteria [38]. This research explores the diversity and composition of microbial communities in the leaf midribs of both HLB-affected and healthy citrus plants. Our research discovered that the Illumina sequencing protocol can be used to evaluate the bacterial endophytes present in plant tissues. The sequencing can be improved with a good choice of primer pair to amplify a longer stretch of the 16S rRNA gene. Our empiric findings illustrate the importance of this platform for accurate and high-resolution microbiota profiling (N90% at species level) of endophytic populations or may be extended to other resources/samples. It was critical to design multiple testing procedures to minimize the bias introduced by host DNA (chloroplast) and chimaera, which were both removed without changing the overall read quality. We have a good likelihood of executing the read sequence efficiently on a specific platform with the Mothur pipeline. This reduced the possibility of read contamination.

The total diversity and complexity of microbiome populations in plant tissues, which include both cultivable and non-cultivable endophytic bacteria, were also exposed using the novel NGS shotgun 16S rRNA gene. Alpha diversity (AD), comprised of species abundance boxplots, species richness curves, and statistical analysis indices, is a common technique for evaluating bacterial diversity within populations [39]. The spreading of bacterial species across tissues and the overall mutual richness is illustrated in this Venn diagram. The Venn diagram (map) of the OTU distribution exposed a colonization pattern of Acinetobacteria 1.41%, Cyanobacteria 28.56%, Firmicutes 1.31%, and Proteobacteria 63.76% of microbes contained in plant leaves were also identified in three samples.

The phylum cyanobacteria were found to be more common in *C. reticulata* Blanco as compared to other phyla. On the contrary, the other two samples presented a greater fraction of Proteobacteria, and few phyla were not observed by culture-based methods, illustrating the importance of NGS. This also led to the fact that these microbes can spread through a variety of channels that penetrate plant tissues [40, 41]. Finally, Proteobacteria, Firmicutes, Actinobacteria, Cyanobacteria, and Bacteriodetes were found to colonize citrus plant leaf tissues, they have been demonstrated to produce useful bioactive chemicals A comparison of bacterial species based on their structure is referred to as beta diversity. As a result, the differences in microbial populations are measured using beta-diversity metrics. A square “distance” or dissimilarity matrix, such as Unweighted Unifrac, was calculated to reflect the contrast among test plant leaves to compare microbial communities between each pair of group samples [42, 43] and Weighted Unifrac distances [44].

At the phylum level, Actinobacteria accounted for 26.47 percent, Cyanobacteria for 2.94 percent, Firmicutes for (23.52%), and Proteobacteria for (47.05%), which was significantly higher than the fraction of other phyla. Though we found 100 genera among them most common were Staphylococcus, Pseudomonas, Lactobacillus, Sphingomonas, Bacillus, Streptomyces, and Pantoea. Bacillus and Lactobacillus, as well as Streptomyces, have previously been found in the roots or leaves of infected (CLas) or infected citrus trees [46–49]. Pantoea, Curtobacterium, and Methylobacterium were also detected in citrus leaves in this analysis. All of these have previously been characterized in terms of bud wood, leaves, and roots [50].

Through studying the PCR products of 16S rDNA sequences covering two specific regions (V3–V4 regions), [45] discovered the diversity of bacterial endophytes from Aloe vera plant leaves, stems, and roots using the NGS by illumina Hi seq technology. The most popular genera identified were Proteobacteria, Firmicutes, Actinobacteria, and Bacteriodetes. This research was identical to the findings of the current study, but we looked for diversity in the V5-V7 region of 16S r RNA. Illumina for next-generation sequencing Hiseq is a relatively new method, with only a limited amount of literature available on it. The discovery of novel bacterial endophytes from citrus illustrates the significance of this study. There has been no comparable work being done with this technique in citrus in other regions of the world, not yet in Pakistan.

### Conclusion

The predominant bacterial groups in the leaf of citrus varieties were Proteobacteria, Actinobacteria, Cyanobacteria, Firmicutes, and Bacteroides, although other groups were commonly found to be less prevalent. Through the culture-dependent method, we find changes in bacterial diversity of endophytes from a citrus leaf but in comparison with an uncultured method, no significant variations existed in relative abundance and diversity of bacteria among taxa from both symptomatic and asymptomatic leaf samples. Some genera such as Staphylococcus, Enterococcus, Enterobacter, Pseudomonas, Bacillus, and Burkholderia were also found in the cultured approach (unpublished data). Although the type of strains has a significant influence on their functional characterization in terms of plant growth-promoting traits rather than their source of isolation either from bulk soil or rhizosphere soil. These genera have been widely found in most of the diversity-related studies of different parts of plants and soils. Some of the isolated strains have great potential to enhance plants growth and they can also be utilized as biocontrol agents against different plant diseases. Finally, this study indicates that these endophytic bacteria may be tested in open field conditions on the same host plants to see whether their biocontrol potential or plant growth-promoting action is successful. Furthermore, their effects on plant physiology could be estimated. We may use these endophytes to produce biofertilizers to replace chemical fertilizers if the same results are obtained from field trials.

## Acknowledgments

The first author acknowledges the financial grant from Higher Education Commission (HEC) Pakistan.

## Notes

### Competing Interest Statement

The authors have declared no competing interest.

